# G2GSnake: A Snakemake workflow for host-pathogen genomic association studies

**DOI:** 10.1101/2023.04.11.536440

**Authors:** Zhi Ming Xu, Olivier Naret, Jacques Fellay

## Abstract

**Summary:** Joint analyses of paired host and pathogen genome sequences have the potential to enhance our understanding of host-pathogen interactions. A systematic approach to conduct such a joint analysis is through a “genome-to-genome” (G2G) association study, which involves testing for associations between all host and pathogen genetic variants. Significant associations reveal host genetic factors that might drive pathogen variation, highlighting biological mechanisms likely to be involved in host control and pathogen escape. Here, we present a Snakemake workflow that allows researchers to conduct G2G studies in a reproducible and scalable manner. In addition, we have developed an intuitive R Shiny application that generates custom summaries of the results, enabling users to derive relevant insights.

**Availability and implementation:** G2GSnake is freely available at: https://github.com/zmx21/G2GSnake under the MIT license.

## Introduction

Hosts and pathogens are involved in a evolutionary battle that involve successive rounds of evolution from both sides (Daugherty and Malik, 2012). Pathogens constantly evolve to escape from host immunity or other control mechanisms, while on a much longer time-scale hosts are also under evolutionary pressure from pathogens. The signatures of this evolutionary battle are reflected on both genomes. Therefore, joint analyses of host and pathogen genomes offer an opportunity to re-capitulate such processes and to identify specific genetic loci involved in host-pathogen interactions.

An effective method to jointly analyze paired host and pathogen genomes the “genome-to-genome” (G2G) approach, which involves searching for significant associations between host and pathogen variants (Fellay and Pedergnana, 2020), reflecting host selection pressure. Such evolutionary conflict often occur between host genetic loci that are involved in pathogen control and pathogen genetic loci that are involved in immune escape. Indeed, G2G studies conducted in viruses such as human Immunodeficiency Virus (HIV) (Bartha et al., 2013), Hepatitis C Virus (HCV) (Ansari et al., 2017; Chaturvedi et al., 2019; Ansari et al., 2019), and Epstein-Barr Virus (EBV) (Rüeger et al., 2021) have uncovered important host-pathogen interaction mechanisms.

Given the generalizability of the G2G approach to many other pathogens, we present here a Snakemake workflow that can be applied to any paired host-pathogen genetic data. We have also developed a R Shiny app that enables users to generate custom queries and visualizations of the results.

## Methods

### Workflow

A summary of the functionalities of the G2GSnake software is shown in Figure 1. In brief, the workflow relies on: 1) pathogen genetic data, either a nucleotide multiple sequence alignment or a amino acid matrix for each gene, 2) host genetic data, in the form of a VCF file, and 3) a sample mapping file that lists sample pairs and additional covariates. Genetic principal components are calculated for both the host and pathogen to correct for stratification. Associations between all host and pathogen variants are then tested under the genome-to-genome (G2G) framework, corrected for principal components and any provided covariates. A directed acyclic graph for a typical workflow is shown in Figure S1. All jobs are executed within a provided docker container to ensure reproducibility and cross-platform compatibility.

**Fig. 1:**
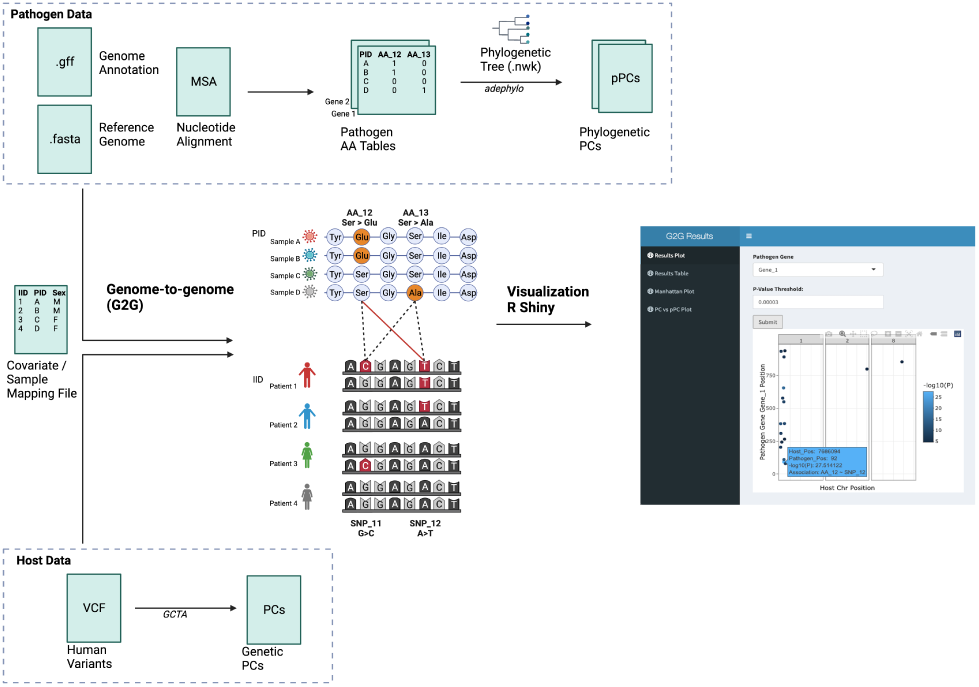
Summary of G2GSnake. The workflow relies on three main input sources 1) pathogen genetic data 2) host genetic data, and 3) a sample mapping file along with additional covariates. Associations between all pathogen and host genetic variants are then tested under the genome-to-genome (G2G) framework. Pathogen and host genetic principal components are included as covariates to correct for stratification. Finally, results can be visualized in the R Shiny app. Created with BioRender.com

### Pathogen Genetic Data

On the pathogen side, a binary matrix, indicating the presence or absence of each amino acid variant is required. This should be separated by gene. We also provide functionalities that derive such matrix from a multiple-sequence nucleotide alignment file (fasta) using custom R scripts and nextalign for translation(Aksamentov et al., 2021). In such an instance, a reference genome (fasta) and annotation (gff) file is also required. Next, given that a phylogenetic tree in newick format is provided, phylogenetic principal components are calculated using the adephylo package in R (Jombart et al., 2010). Alternatively, standard principal components can be calculated. Finally, pathogen variants are filtered based on user-specified allele frequency and missingness thresholds.

### Host Genetic Data

On the host side, a VCF file is required. Quality control procedures are based on user-defined thresholds and includes filtering based on missingness, minor allele frequency, and deviation from Hardy-Weinberg equilibrium. Principal components are then calculated using GCTA (Yang et al., 2011).

### Genome-to-genome study

The software relies on existing tools developed for genome-wide association studies (GWAS). A case-control GWAS is run for each pathogen variant, treating it as an outcome. Users can choose to use PLINK (logistic regression) (Chang et al., 2015) or SAIGE (generalized linear mixed models) (Zhou et al., 2018). SAIGE can provide more reliable estimates for rare pathogen variants. However, for very large pathogen genomes, using SAIGE could be computationally infeasible.

### R Shiny App

To visualize and summarize results from the G2G study, we developed a R Shiny app. The app is launched from a Docker container to ensure cross-platform compatibility. The app provides four main functions: 1) A results plot which displays all G2G associations in the specified pathogen gene and under a specified p-value threshold, 2) A results table which displays summary statistics of all G2G associations under a specified p-value threshold, 3) A manhattan or QQ plot for a specified pathogen variant or all variants, and 4) A correlation plot between host and pathogen principal components.

## Results

To test the validity of our software, we generated a simulated host-pathogen genetic dataset that includes variants that were population stratified but also true associations between certain pairs of host and pathogen variants. We selected parameters based on a previous G2G simulation study (Naret et al., 2018), but reduced both the sample size and number of variants by a factor of 10 for computational reasons. In summary, the simulation study included 500 paired samples with 5030 host variants and 40 pathogen variants. Pathogens and hosts were each divided into two sub-populations. 5010 host variants were not associated with any pathogen variants, with 4000 of them not stratified, 1000 of them stratified across host sub-populations, and 10 of them stratified across both host and pathogen sub-populations. 20 host variants were associated with 20 of the pathogen variants, with 10 pairs not stratified and the other 10 pairs stratified across both host and pathogen sub-populations. The remaining 20 pathogen variants were not associated with any host variants, with 10 of them stratified across pathogen sub-populations and the other 10 stratified across both pathogen and host sub-populations.

We utilized the top 5 host and pathogen principal components to correct for stratification. Figure S6 summarizes the associations detected by G2GSnake when applied to the simulated data. Pairs of host and pathogens variants could be classified into three main categories: 1) true G2G associated pairs 2) true G2G associated pairs but stratified across host and/or pathogen sub-populations, or 3) not associated pairs. As expected, associations between true G2G associated pairs were detected by the software as strongly associated. For stratified G2G associated pairs, the detected association was less significant given that principal components cannot fully separate true signal from stratification. For pairs that were not associated, the detected association was negligible. Few non-associated pairs were detected as weakly associated likely due to residual stratification that principal components were not able to capture.

## Conclusion

We introduce a scalable and reproducible Snakemake workflow that allows researchers to jointly analyze paired host-pathogen genetic data under the genome-to-genome (G2G) framework. This tool would allow G2G studies to be systematically conducted and interpreted for any pathogen of interest. Docker containers are provided for the workflow itself and the R Shiny app to ensure cross-platform compatibility.

## Supplementary Materials

**Fig. S1:**
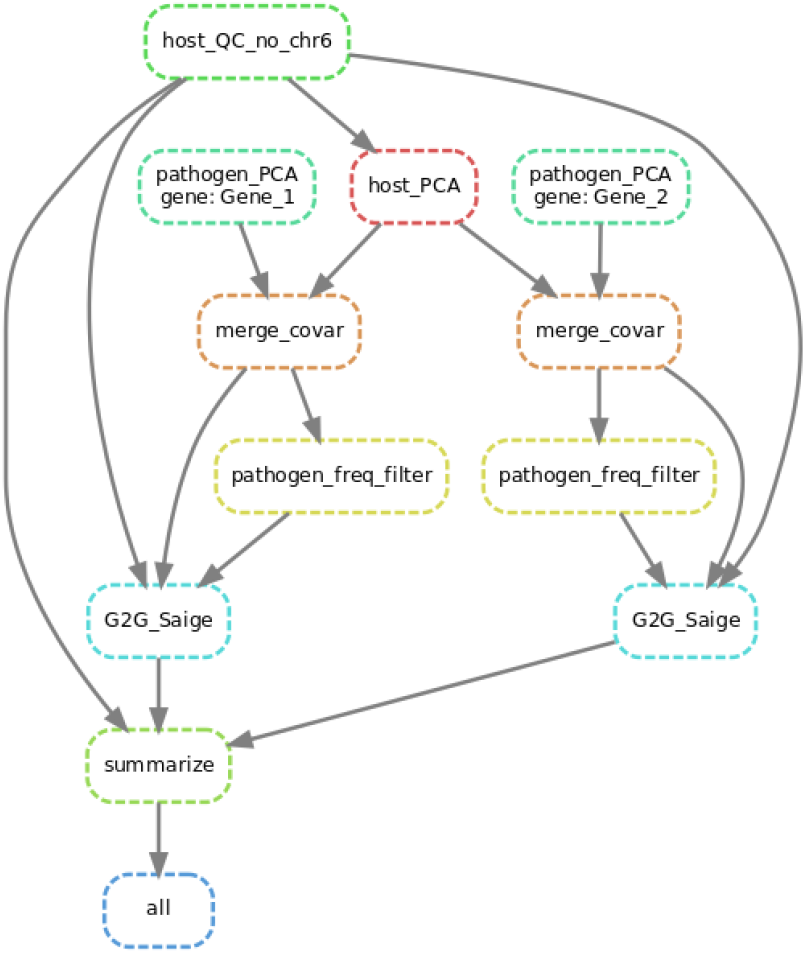
Directed acyclic graph of snakemake rules. The workflow steps can be summarized as: 1) QC on host genetic data 2) Calculation of host PCA 3) Calculation of pathogen PCA for each pathogen gene 4) Variant frequency filter on pathogen data, separately for each gene 5) Genome-to-Genome study for each gene 6) Write out summary statistics and results files that be read by the R Shiny app

**Fig. S2:**
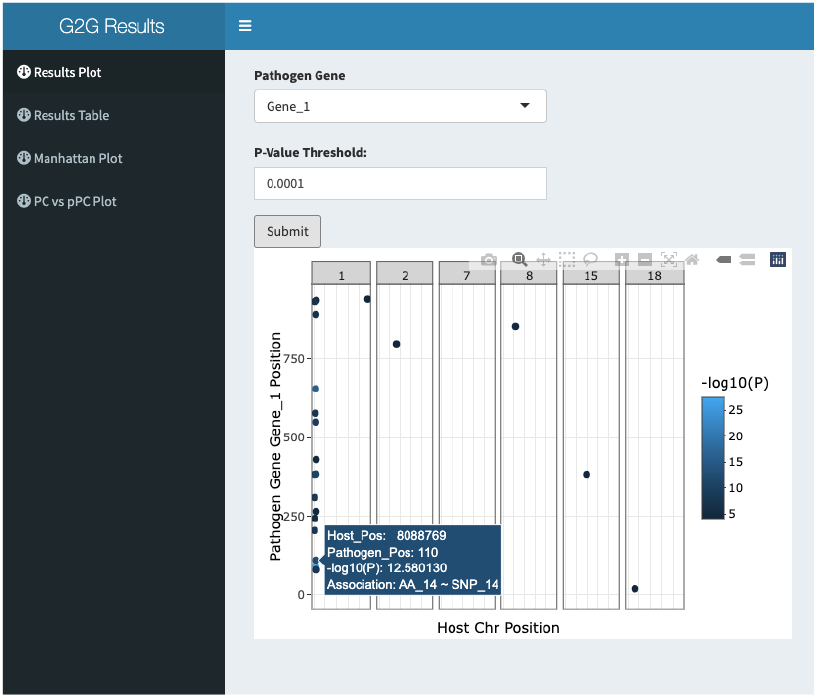
Shiny App - Results Plot.

**Fig. S3:**
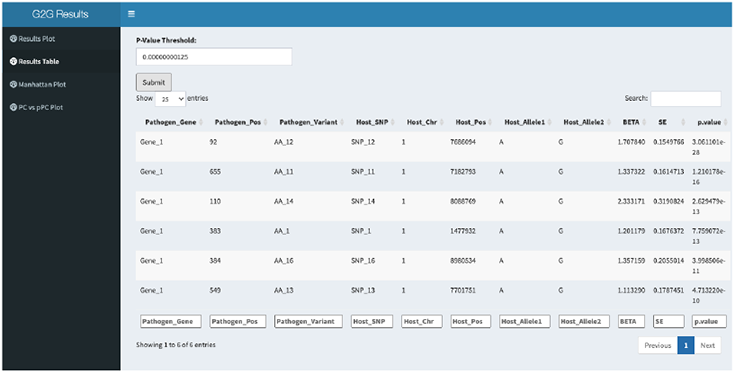
Shiny App - Results Table.

**Fig. S4:**
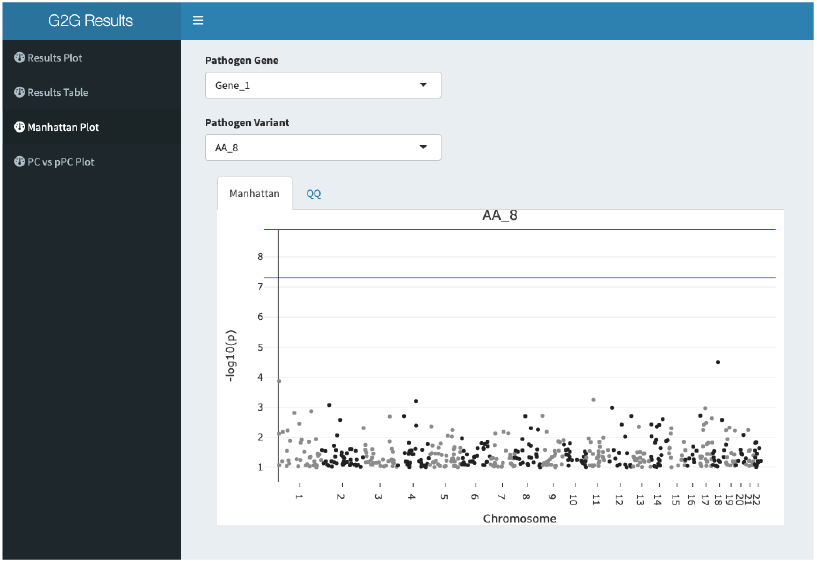
Shiny App - Manhattan Plot.

**Fig. S5:**
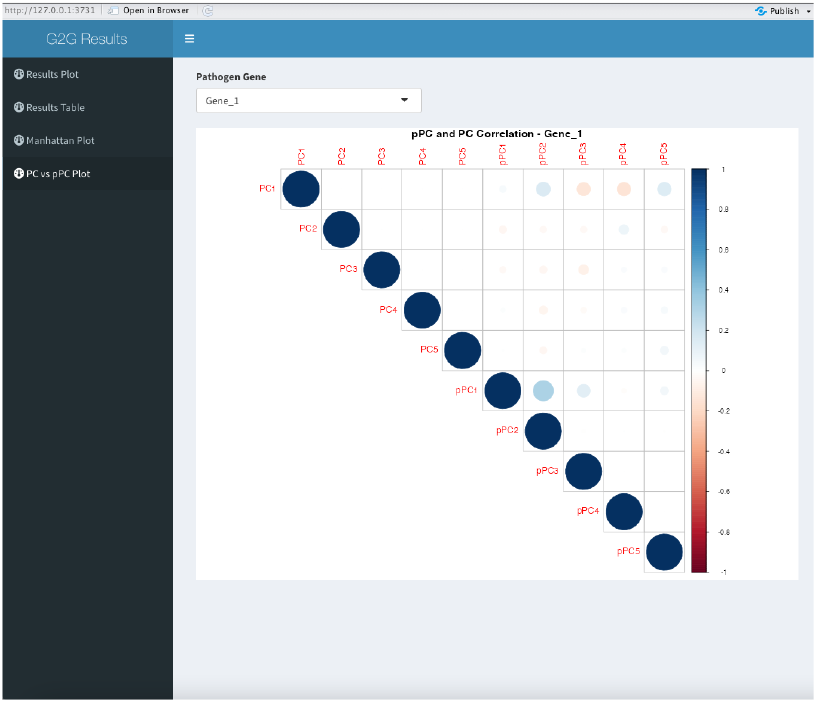
Shiny App - PC vs pPC Plot.

**Fig. S6:**
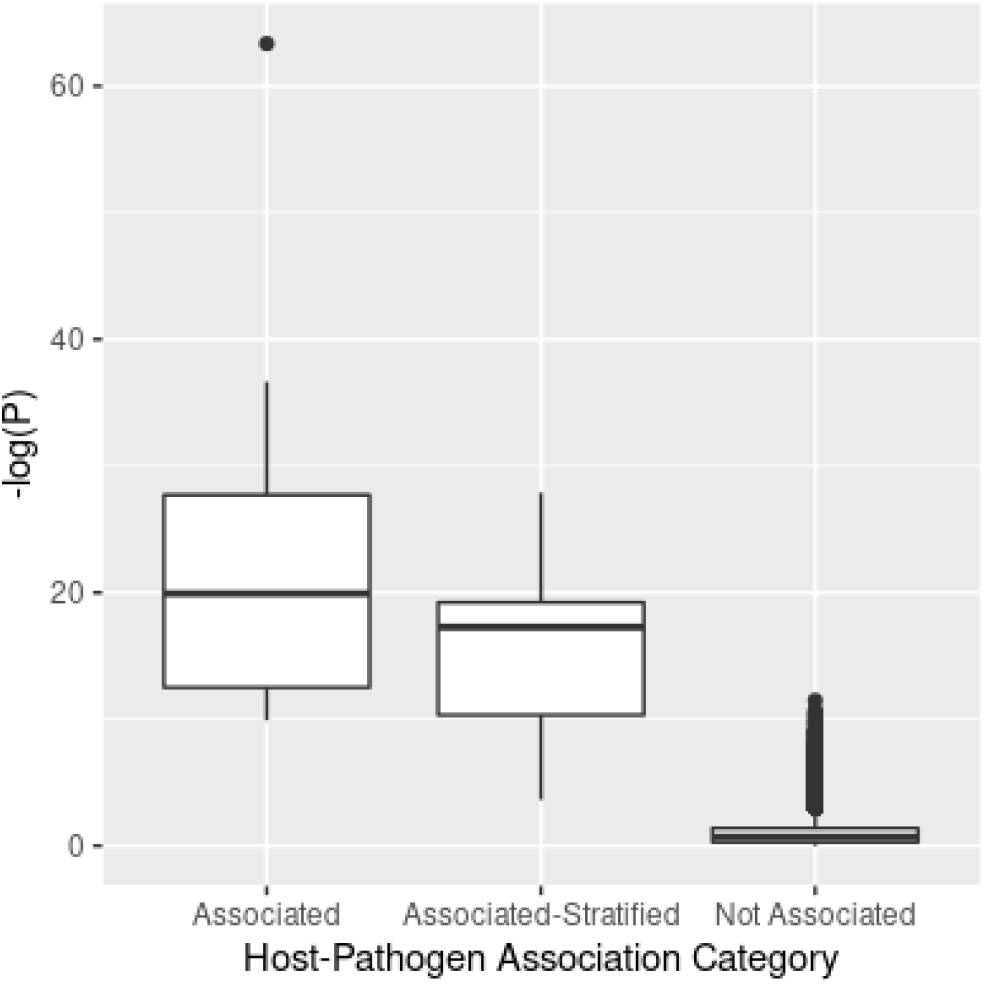
Simulation Results. Y-axis indicates -log(P) of G2G associations reported by G2GSnake. X-axis indicates simulated category of pairs of host and pathogens variants.

