## Supplementary Materials for "G2GSnake: A Snakemake workflow for host-pathogen genomic association studies"

Fig. S1: **Directed acyclic graph of snakemake rules** The workflow steps can be summarized as: 1) QC on host genetic data 2) Calculation of host PCA 3) Calculation of pathogen PCA for each pathogen gene 4) Variant frequency filter on pathogen data, separately for each gene 5) Genome-to-Genome study for each gene 6) Write out summary statistics and results files that be read by the R Shiny app

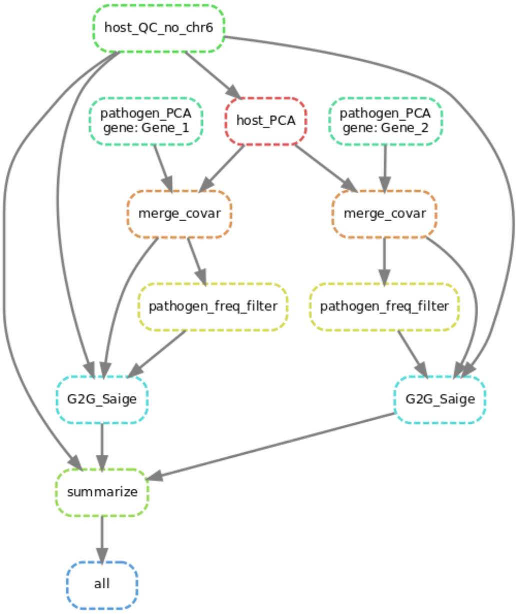


Fig. S2: **Shiny App - Results Plot**

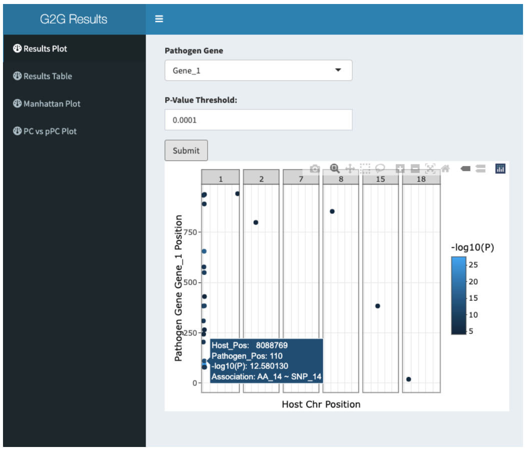


Fig. S3: **Shiny App - Results Table**

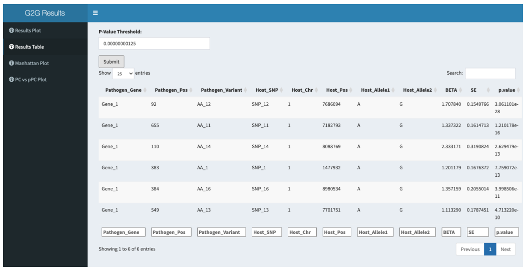


Fig. S4: **Shiny App - Manhattan Plot**

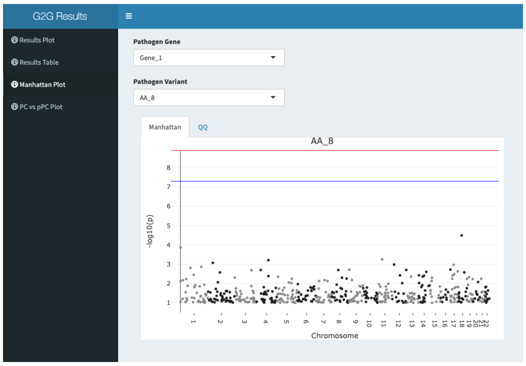


Fig. S5: **Shiny App - PC vs pPC Plot**

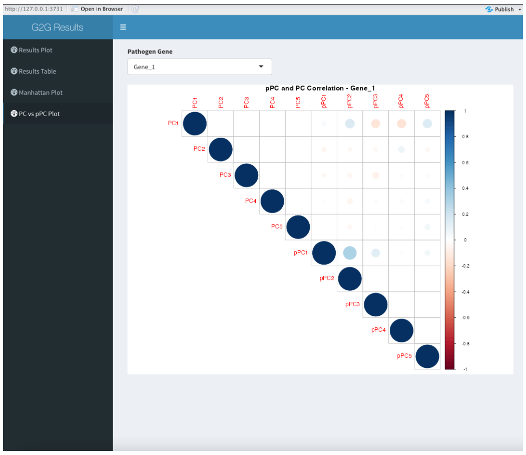

Fig. S6: **Simulation Results** Y-axis indicates -log(P) of G2G associations reported by G2GSnake. X-axis indicates simulated category of pairs of host and pathogens variants.

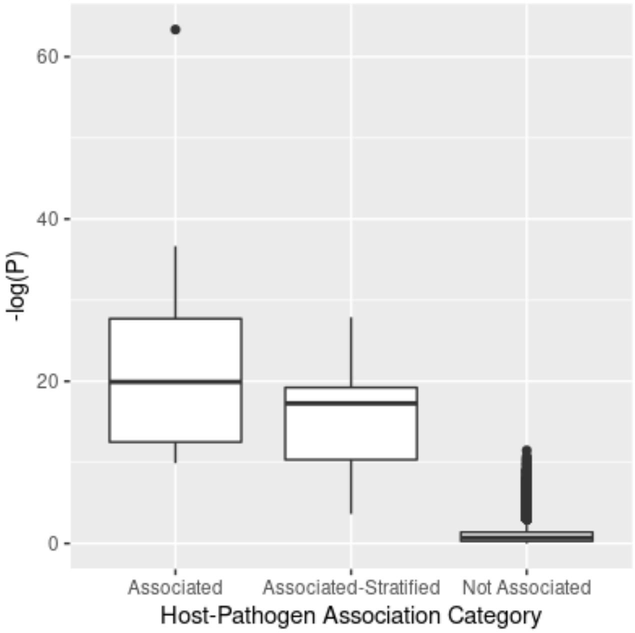
